# Unravelling mechanisms of meropenem induced persistence facilitates identification of GRAS compounds with anti-persister activity against *Acinetobacter baumannii*

**DOI:** 10.1101/2020.07.31.231936

**Authors:** Timsy Bhando, Ananth Casius, Siva R. Uppalapati, Ranjana Pathania

## Abstract

*Acinetobacter baumannii* is recognized as one of the “critical” pathogens by the World Health Organisation (WHO) due to its unprecedented ability to acquire resistance genes and undergo genetic modifications. Carbapenem classes of antibiotics are considered as the “drugs of choice” against *A*. *baumannii* infections, although increasing incidence of carbapenem resistant isolates have greatly limited their efficacy in clinical settings. Nonetheless, the phenomenon of multi-drug tolerance or persistence exhibited by *A*. *baumannii* has further led to therapeutic failure of carbapenems against chronic and recurring infections. Exploring the underlying mechanisms of persistence hosted by the nosocomial pathogen, *A*. *baumannii* can facilitate the development of effective anti-persister strategies against them. Accordingly, this study investigates the characteristics and mechanisms responsible for meropenem induced persistence in *A. baumannii.* Furthermore, it describes the adaptation of a screening strategy for identification of potent anti-persister compounds that cumulatively act by targeting the *A. baumannii* membrane, inhibiting antibiotic efflux and inducing oxidative stress mediated killing. The screen identified the phytochemical compound, thymol to display excellent activity against persisters of mechanically distinct antibiotics. While meropenem exposed *A. baumannii* persisters exhibited multi-drug tolerance and indicated the ability to enter a Viable But Non Culturable (VBNC) state, thymol efficiently eradicated all persister cells, irrespective of their culturability. Thymol exhibited no propensity for resistance generation and also inhibited persisters of other Gram-negative pathogens, *Pseudomonas aeruginosa* and *Klebsiella pneumoniae.* Collectively, our results establish thymol to have immense potential to act either alone or as an adjunct in combination therapies against persistent infections.

**IMPORTANCE:** Apart from the global catastrophe of antibiotic resistance, the phenomenon of “antibiotic tolerance” exhibited by a subpopulation of bacterial cells known as “persisters” ensue a major clinical threat. Eradication of the persister populations holds extreme importance for an improved long-term recovery from chronic and recurring bacterial infections. This study addresses the problem of antibiotic persistence prevailing in clinics and investigates its associated mechanisms in the nosocomial pathogen, *Acinetobacter baumannii* in reference to the antibiotic meropenem. It further describes the use of a mechanism-based screening approach for the identification of potent multi-targeting anti-persister compounds, thereby leading to the identification of GRAS (Generally Regarded As Safe) molecules exhibiting promising activity against *A*. *baumannii* persisters. This strategy can further be utilized for repurposing of FDA approved drugs or other available compound libraries, in order to identify novel anti-persister compounds.

## INTRODUCTION

Antibiotic resistance is one of the major global health concerns. Imprudent and indiscriminate use of antibiotics has led to the emergence of multidrug resistant (MDR) and extensively drug resistant (XDR) bacterial strains causing more than 23,000 deaths per year in the United States (1). The “ESKAPE” pathogens (*Enterococcus faecium, Staphylococcus aureus, Klebsiella pneumoniae, Acinetobacter baumanii, Pseudomonas aeruginosa*, and *Enterobacter* species) are known to cause a plethora of untreatable infections ranging from septicemia, pneumonia to urinary tract infections (UTIs). Importantly, Gram-negative bacteria make up four of the six so-called “ESKAPE” pathogens and are particularly difficult to treat owing to the presence of multidrug efflux pumps and complex envelope architecture (2). With the ever-increasing rise in “global resistome” and significant decline in novel antibiotic discovery, there is an urgent need for alternative strategies to tackle the current predicament.

*A. baumannii* is one of the most troublesome Gram-negative ESKAPE pathogens, responsible for causing a wide range of hospital acquired infections such as recurring UTIs, secondary meningitis, wound infections and ventilator-associated pneumonia (VAP) (3). The Infectious Disease Society of America (IDSA) has declared *A. baumannii* as one of the ‘red alert’ pathogens, thus highlighting the threat it poses to the healthcare systems (4). In addition to the organism’s high levels of intrinsic resistance, *A. baumannii* exhibits extreme genetic flexibility and remarkable ability to acquire resistance determinants in response to selective environmental pressure (5). The uncanny ability of this bacterium to resist desiccation, survive harsh environmental conditions and form biofilms, both on medical devices and biological surfaces qualifies it to be a dreadful nosocomial pathogen (3). Alternatively, *A. baumannii* is also known to enter a physiological state where it becomes refractory or tolerant to the action of antibiotics (6,7). This phenomenon, referred to as ‘persistence”, is the major cause for antibiotic failure and relapsing infections in clinical settings (8). Eradication of the persister populations in chronic ailments holds extreme importance for an improved long-term recovery from recalcitrant infections. Meropenem, tigecycline, rifampicin and polymyxin B are commonly administered against *A. baumannii* infections in hospital settings, although increasing incidence of both resistant and persistent isolates have limited their efficacy (7, 9–17). Therefore, screening for novel agents that can inhibit antibiotic tolerant cells presents an unusual challenge and can provide a successful alternative in the fight against recurring *A. baumannii* infections (18–20). Since most of the clinically important antibiotics act on DNA, RNA or protein targets that exhibit quiescence during “persistent” infections; membrane-active agents have emerged as an important new means of eradicating recalcitrant dormant bacteria (21, 22). Furthermore, recent studies have reported the crucial role of reactive oxygen species (ROS), membrane potential and efflux mechanisms for drug tolerance in persistent populations (23, 24).

In this study, we examined the ability of *A. baumannii* to produce antibiotic tolerant persisters to clinically relevant antibiotics from different classes, both in planktonic phase as well as biofilms. Carbapenem class of antibiotics have long been the mainstay of therapy against *A. baumannii* infections (25). Therefore, meropenem, a commonly used carbapenem remained the focus for our study. Here, we preliminarily investigated the characteristics and mechanisms of meropenem tolerance in *A. baumannii*. Furthermore, a library of molecules encompassing the GRAS (Generally Regarded As Safe) status was screened for their “anti-persister” potential; based on their ability to target the bacterial membrane, cause increased ROS production and inhibit the proton motive force (PMF) and multidrug efflux pumps. The screen identified thymol to possess excellent inhibitory activity against *A. baumannii* persisters. To our knowledge, this is the first study that utilizes a mechanism-based screen to identify novel anti-persister compounds against *A. baumannii*. Importantly, this study for the first time reports the discovery of GRAS status natural compounds demonstrating excellent inhibitory activity against *A. baumannii* persisters. These findings can have vital implications for the improved treatment of recalcitrant *A. baumannii* infections in the clinics.

## RESULTS

### *A. baumannii* AYE formed persisters in response to high concentrations of Meropenem, tigecycline, rifampicin and polymyxin B

The Minimum Inhibitory Concentration (MIC) of meropenem, rifampicin, tigecycline and polymyxin B against *A. baumannii* AYE were determined to 0.5 μg/mL, 4 μg/mL, 0.25 μg/mL and 0.5 μg/mL respectively; as per CLSI guidelines. Antibiotic tolerant *A. baumannii* persisters were isolated by exposing stationary phase (SP) *A. baumannii* AYE cells to high concentrations of meropenem (100X MIC), tigecycline (50X MIC), rifampicin (50X MIC) and polymyxin B (40X MIC) for 12h and the number of surviving bacteria were determined (26–28). All four antibiotics demonstrated the presence of significant fraction of surviving persisters (Fig. 1A). Persistence frequencies were calculated w.r.t untreated SP cells and meropenem displayed the maximum frequency of 0.42±0.01%. Rifampicin, tigecycline and polymyxin B also showed high persistence frequencies of 0.29±0.01%, 0.297±0.03% and 0.3±0.02%, respectively. The susceptibility of isolated persisters against respective antibiotics were re-determined by broth microdilution, to confirm non-acquisition of resistance mechanisms (28). Isolated persisters were subsequently regrown into fresh medium and passaged up to three generations followed by determination of MIC and persistence frequency, after each passage. Isolated fractions showed similar antibiotic susceptibility and survival frequencies, thus confirming the isolation of true persister cells.

**FIG 1.**
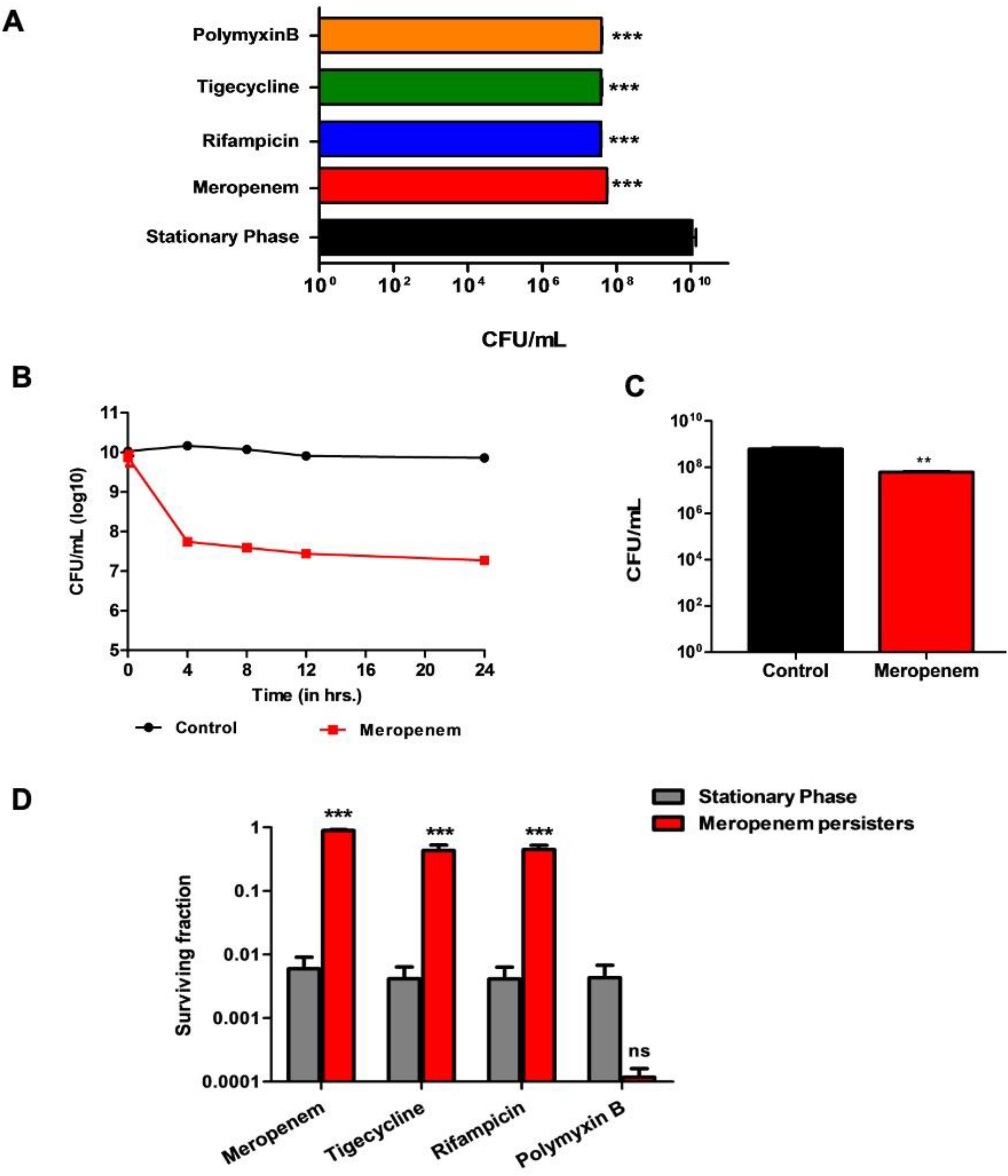
(A) *A. baumannii* AYE formed persister in response to Meropenem (100X MIC), Tigecycline (50X MIC), Rifampicin (50X MIC) and Polymyxin B (40X MIC). Data are means±SEM. P values determined by one-way ANOVA and Tukey’s multiple comparison test (B) Meropenem induced *A. baumannii* AYE persisters reveal a biphasic killing pattern. (C) *A. baumannii* AYE biofilms exhibit tolerance to meropenem. P values were determined by aired t-test (two-tailed) (D) Meropenem induced *A. baumannii* persisters exhibit multi-drug tolerance. Gray bars denote fraction of stationary phase cells that survive exposure to respective antibiotics. Red bars denote fraction of meropenem persisters that survive successive exposure to indicated antibiotics. Significance was calculated using two-way ANOVA and Bonferroni post hoc test test (**, P<0.01; ***, P<0.0001; ns, non-significant).

### Meropenem induced *A. baumannii* AYE persisters revealed a biphasic killing pattern

The state of bacterial persistence or dormancy is characterized by typical biphasic killing pattern in the presence of bactericidal concentration of antibiotics (8). Biphasic curves display an initial drastic killing of bulk of bacterial population followed by occurrence of a plateau phase where only the surviving persister fractions sustain. Biphasic killing patterns in several bacteria are both time and concentration dependent (29). However, isolates of *Acinetobacter calcoaceticus-baumannii* (ACB) complex are reported to exhibit high persister frequencies independent of antibiotic concentration, thereby complicating treatment efficacy (17). We exposed SP cells of *A. baumannii* AYE to meropenem (100X MIC) and observed a rapid decrease in viable count (2.36±0.01 log) up to 4h, followed by relatively constant CFU until 24h (Fig. 1B). The biphasic pattern obtained thus revealed the presence of meropenem tolerant *A. baumannii* persisters. Persisters were further re-inoculated into fresh medium and subsequently exposed to 100X MIC of meropenem. Re-isolated persisters displayed similar antibiotic susceptibility, persistence frequency and biphasic killing; thereby confirming non-heritability of resistance.

### Meropenem induced persister formation under biofilm conditions

*A. baumannii* exhibits a remarkable ability to adhere and form biofilms on both biotic and abiotic surfaces such as site of wound infections, hospital surfaces and medical devices (30, 31) Biofilm communities are known to contain persister subpopulations which are responsible for their recalcitrant and relapsing nature (32). *A. baumannii* AYE biofilms (48h old) grown on 96-well plates were exposed to meropenem (at 100X MIC) and persister levels were detected after 12h. A significant fraction of biofilm associated cells on polystyrene surface (7.78±0.02 log CFU/ml) survived exposure to bactericidal concentration of meropenem, yielding persister frequency of 0.1±0.02% (Fig. 1C).

### Meropenem persisters of *A. baumannii* exhibit multidrug tolerance

Since persisters are dormant bacterial cells that exhibit inactive targets, these are known to display tolerance to multiple antibiotics (33). Hence, we sought to investigate the tolerance of meropenem persisters of *A. baumannii* AYE towards antibiotics acting on varying cellular targets. Cross-tolerance was measured by exposing isolated meropenem persisters to rifampicin (50X MIC), tigecycline (50X MIC) and polymyxin B (20X MIC) for 12h. The fraction of meropenem persisters that survived exposure to test antibiotics was compared to the surviving fraction of SP cells (34). As shown in Fig. 1D, significant fraction of meropenem persisters survived exposure to tigecycline and rifampicin, thus exhibiting a high degree of cross-tolerance and indicating the presence of overlapping persistence mechanisms. On the contrary, polymyxin B inhibited meropenem persisters and extremely low fractions of surviving bacteria were obtained. This indicated that the mechanisms responsible for persistence to meropenem and polymyxin were exclusive to each other and could not occur simultaneously (35). A similar observation was reported in a previous study, where meropenem-polymyxin combination inhibited *A. baumannii* persisters (36).

### Meropenem induced *A. baumannii* persisters exhibit altered morphology

The lipophilic membrane staining dye, FM 4-64FX was used to assess the morphology of meropenem induced *A. baumannii* persisters. While most cells in the isolated persister fractions exhibited the typical cocci shape (1-2 μm), few cells were observed to display an elongation phenotype with cell size in the range 3-6 μm (Fig.2A). These cells reverted into the untreated *A. baumannii* morphology in absence of meropenem. Recently, stationary phase cultures of colistin-heteroresistant *A. baumannii* strains were reported to contain elongated rod-like cells by unknown mechanisms (37). The occurrence of elongated cells in a fraction of meropenem persisters is intriguing and warrants further investigation to decipher underlying mechanisms.

**FIG 2.**
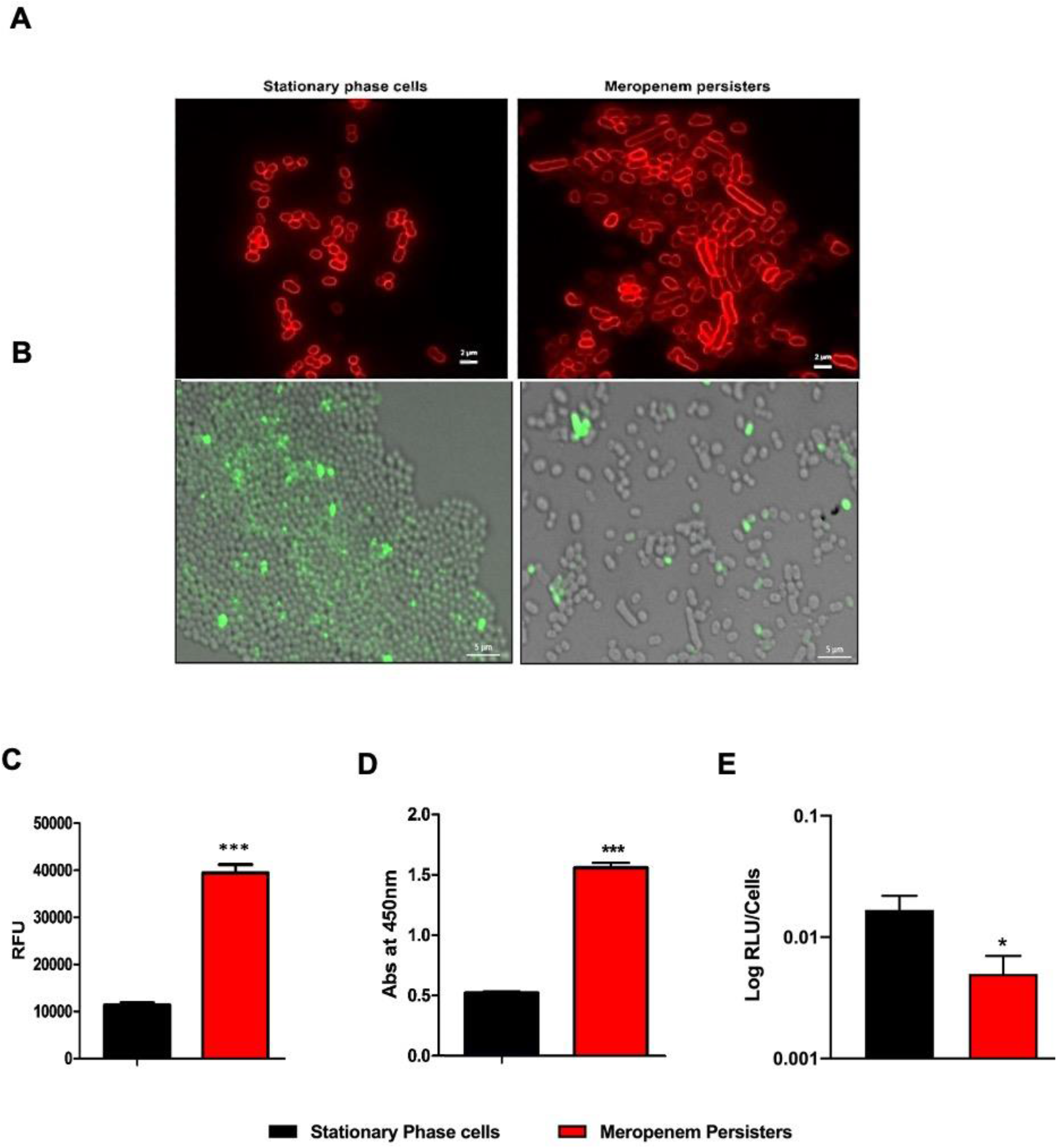
(A) Meropenem induced *A. baumannii* AYE persisters exhibit altered morphology. Cells were stained with FM 4-64FX. Scale bar represents 2 μm. (B) Meropenem induced *A. baumannii* persisters exhibit decreased accumulation of BOCILLIN, in comparison to stationary phase cells. Overlay of brightfield and fluorescent image is shown. Scale bar represents 5 μm. Meropenem induced *A. baumannii* persisters exhibit (C) Membrane depolarization and (D) High respiratory activity in XTT assay and (E) ATP Levels, in comparison to stationary phase cells. Data are means±SEM. P values were determined by unpaired t-test (two-tailed) (*, P<0.05; **, P<0.01; ***, P<0.0001).

### Meropenem persisters exhibit decreased antibiotic accumulation

Membrane permeability in bacteria is the net effect of efflux pumps and porin proteins which are responsible for antibiotic uptake (24). Recent studies have reported the crucial contribution of active efflux mechanisms towards bacterial persistence in *E. coli* and *Mycobacterium* species (24, 38). Fluorescence based single cell microscopic studies in *E.coli* revealed lesser antibiotic accumulation within persister cells, thereby indicating increased efflux or decreased membrane permeability (24). Therefore, the accumulation of fluorescent analog of penicillin, BOCILLIN™ FL was studied in persister populations, in comparison to untreated SP *A. baumannii* cells. Interestingly, meropenem persisters were observed to display reduced fluorescence in comparison to SP cells, indicating enhanced membrane impermeability or increased efflux activity, in order to survive exposure to lethal antibiotic doses (Fig. 2B and Figure S1, S3).

Since antibiotic accumulation in Gram-negative bacteria is the net result of interplay of two crucial factors, membrane permeability and efflux activity. Therefore, we used deletion mutants in *A. baumannii* ATCC 17978 of genes encoding an RND efflux pump, AdeIJK and porin protein, Omp33 to study their role in antibiotic induced persistence. AdeIJK is one of the major constitutively expressed efflux pumps in *A. baumannii*, which exhibits a broad substrate specificity (39). ATCC 17978 Δ*adeIJK* mutant showed a decreased ability to survive meropenem exposure in comparison to wild type (Fig S3A). Furthermore, AdeIJK expression in the complemented strain displayed roughly 100 fold more persister levels than that of wild type cells, hence recovering the low persister frequency in the efflux mutant. Reduced expression of several porins such as Omp33-36, CarO, Omp22-33, Omp37 etc. have also been associated with carbapenem resistance in *A. baumannii* (40, 41). The Δ*omp33* mutant displayed higher survival than wild-type ATCC 17978 over a gradient of meropenem concentrations, due to the unavailability of porin for antibiotic uptake (Fig S3B). Concomitantly, the Δ*adeIJK* mutant showed a higher BOCILLIN fluorescence in contrast to Δ*omp33* which exhibited decreased accumulation due to lack of porin (Fig S3C).

### Meropenem induced persisters exhibit membrane depolarization

Considering the importance of membrane permeability in meropenem persistence, we set out to determine the role of various envelope stressors. Membrane potential (MP) refers to the presence of an electrochemical gradient across the inner membrane of bacterial cells leading to generation of PMF that aids proton transfer across membrane, thereby producing ATP (42). Bacterial MP and intracellular ATP levels are known to influence the phenomenon of persistence such that decreased PMF leads to a characteristic low metabolic activity in dormant persister cells. Several Toxin-Antitoxin (TA) systems have been implicated to modulate the MP leading to persistence (43). The membrane potential of meropenem persisters was evaluated using the anionic lipophilic dye, DiBAC_4_(3), that enters depolarized cells and causes enhanced fluorescence intensity (44). Meropenem induced persisters of *A. baumanii* AYE exhibited a significant increase in fluorescence, thus indicating membrane depolarization and a decreased PMF, which is also a characteristic of persistence in *E. coli* cells (Fig. 2C) (44).

### Meropenem treated cultures of *A. baumannii* indicate the presence of VBNC bacteria

As discussed previously, persisters represent a subpopulation that exhibits a state of dormancy characterized by low metabolic activity (45). Hence, we evaluated the respiratory activity of meropenem induced persisters by the XTT assay (2,3-bis[2-methyloxy-4-nitro-5-sulfophenyl]-2H-tetrazolium-5-car-boxanilide). The assay is based on the reduction of the tetrazolium dye, XTT into a water soluble formazan product, which correlates to respiratory activity and levels of NADH within bacterial cell (46). Metabolic activity of SP *A. baumannii* AYE was compared with meropenem persisters and the numbers of CFU in the two conditions were normalized prior to the experiment (Fig S2A). Surprisingly, the assay revealed meropenem persisters to exhibit significantly high respiratory activity in comparison to SP cells (Fig. 2D). This observation indicated the presence of VBNCs, other than persisters in meropenem treated cultures, which obscured the measurement of persister metabolism. Beta-lactam treated cultures of *E. coli* are also reported to contain both persister cells and VBNCs (47).

This observation was further validated by normalizing the number of viable bacteria in both SP cells and persister fractions using flow cytometry and followed by determination of intracellular ATP levels. It was interesting to note that on the basis of culturable CFU counts, meropenem persisters displayed high luminescence i.e. increased ATP levels, well in accordance with the result from XTT assay (Fig S2B). However, when the viable cell counts were normalized via flow cytometry analysis, meropenem persisters exhibited low intracellular ATP, thus demonstrating reduced energy pools (Fig 2E, S2C).

### Meropenem induced *A. baumannii* persisters exhibit similar characteristics, irrespective of the method of isolation

With the recent surge of research on antibiotic persisters and their clinical relevance, published studies have reported varying protocols for persister isolation. In order to overcome the shortcomings associated with *in vitro* analysis of persistence and to have a consensus, Balaban *et al*. outlined the definitions and procedures for persister isolation (28). Having established the underlying mechanisms in meropenem persistence with one protocol, we next sought to assess the characteristics of meropenem persisters in *A. baumannii* isolated following another common method of isolation. In this procedure, 12h old culture of *A. baumannii* AYE was diluted (1:10) and exposed to meropenem (at 10X MIC), as described by Pu *et al* (24). Persister cells thus isolated displayed a biphasic killing pattern, but a rounded phenotype (a.k.a spheroplasts), which reverted into the untreated cell morphology, in the absence of meropenem (Fig 3A). This observation was very much in accordance with a recent study which reported meropenem tolerance in *Enterobacter spp.* and *Klebsiella spp.* to be spheroplast mediated (48). Persisters isolated by diluting stationary phase cells displayed reduced antibiotic accumulation, depolarized membranes and occurrence of VBNCs (Fig 3B, 3C, 3D and S2D).

**FIG 3.**
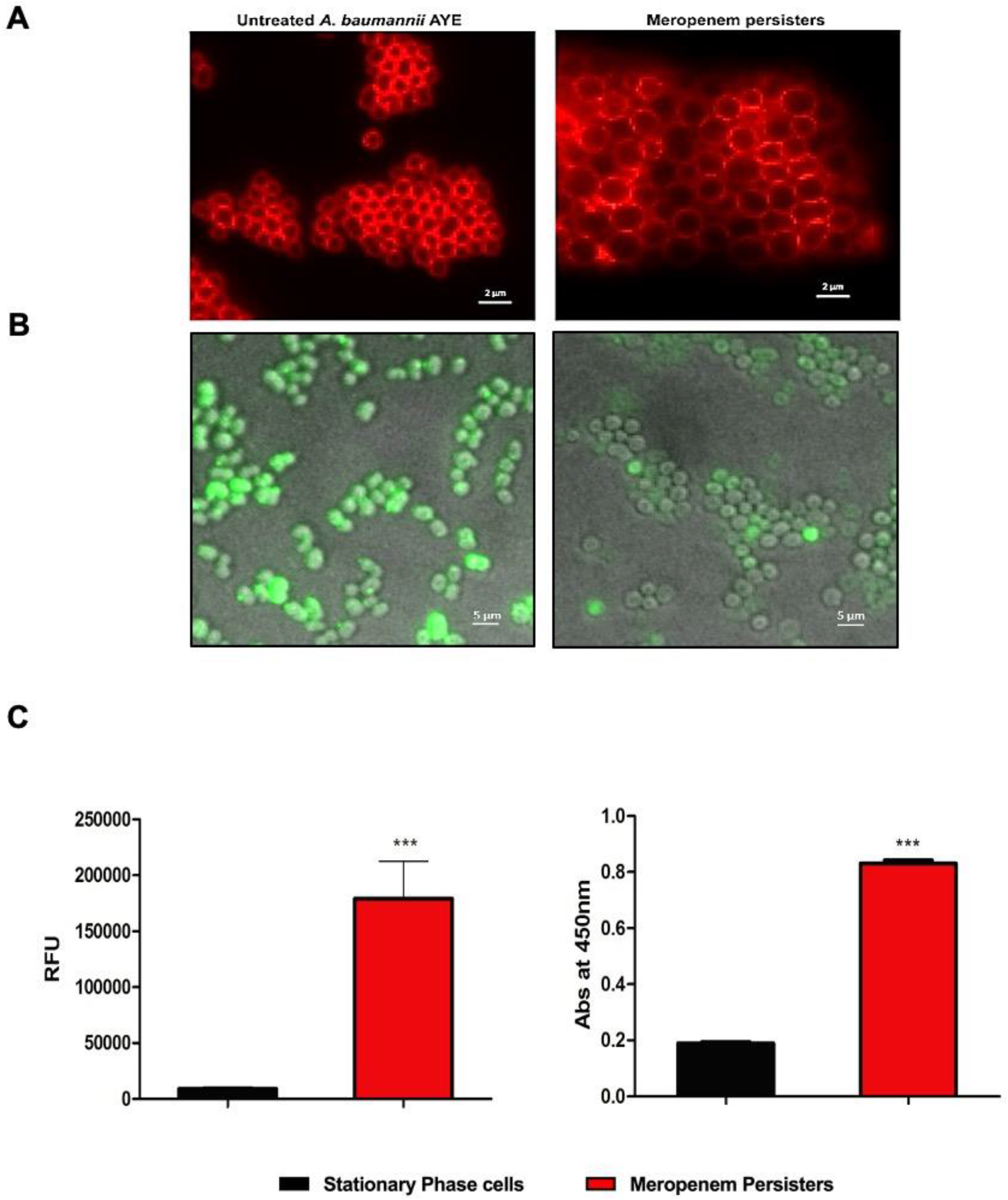
(A) Meropenem induced *A. baumannii* persisters isolated by diluting stationary phase cells exhibit rounded phenotype. Cells were fluorescently stained with FM 4-64FX. Scale bar represents 2 μm. *A.baumannii* persisters displayed (B) Reduced accumulation of BOCILLIN. Overlay of brightfield and fluorescent image is shown. Scale bar is 5 μm (C) Membrane depolarisation and (D) High respiratory activity in the XTT assay. Data are means±SEM. P values were determined by unpaired t-test (two-tailed). (*, P<0.05; **, P<0.01; ***, 0.0001).

### Mechanism based screening for identification of potential anti-persister compounds

Based upon the observed characteristics of *A. baumannii* persisters as well as previous published reports, a mechanism-based screen for the screening and discovery of potent anti-persister compound against *A. baumanni* persisters was devised. An in-house collection of GRAS status small molecules was screened for their ability to target three major mechanisms of bacterial persistence i.e. membrane barrier, efflux pumps and oxidative stress. The MIC of the compounds against *A. baumannii* AYE was initially determined using broth microdilution, as per the CLSI guidelines (Table S1) (49). The compounds carvacrol, thymol, cinnamaldehyde, clove oil and eugenol were observed to possess antibacterial activity against *A. baumannii* AYE (MICs in the range 128-512 μg/mL). Other compounds e.g. linalool and origanum oil displayed higher MIC (≥1 mg/ml) and were therefore excluded from further assays.

The most common strategy to inhibit persisters is through extensive damage of the bacterial membrane (50). Compounds were assessed for their ability to cause membrane damage at sub-inhibitory concentrations by aid of dyes N-phenyl-1-napthylamine (NPN) and SYTOX Orange, which are commonly used to probe the outer and inner membrane damaging potential respectively (51). The outer membrane permeability assay revealed thymol, carvacrol and linalool to exhibit maximum potential to permeabilize the *A. baumannii* outer membrane (Fig. S4A). Likewise, eugenol, thymol and carvacrol displayed enhanced ability to compromise the inner membrane (Fig. S4B).

Bacterial cells produce ROS, namely superoxide and hydrogen peroxide, in response to various stresses and as a metabolic by-product (52). In order to restore the cellular damage inflicted upon by ROS, bacteria display an oxidative stress response, by virtue of antioxidant enzymes. Active suppression of oxidative stress and decreased ROS production is a known mechanism for antibiotic tolerance in bacteria (53). Hence, screening for molecules with pro-oxidant property could be an effective strategy to target persisters (50). Dichloro-dihydro-fluorescein diacetate (DCFH-DA) was used to evaluate the potential of compounds for enhanced ROS production (54). The compounds linalool, thymol and carvacrol acted as pro-oxidants against *A. baumannii* cells (Fig. S4C).

Active efflux mechanisms have been established to contribute significantly to antibiotic tolerance in bacteria (24).Therefore, administration of efflux pump inhibitors (EPI) in combination with antibiotics is an important strategy to target the persister phenotype. Hence, the aforementioned molecules were further screened for their potential to inhibit efflux of Ethidium Bromide (EtBr), a broad substrate inhibitor for most efflux pumps in Gram negative bacteria (55). The assay revealed thymol to be the most potent EPI, followed by eugenol, clove oil, linalool and carvacrol (Fig. S4D). Eugenol and cinnamaldehyde were recently reported to act as EPI in *A. baumannii* against beta-lactam antibiotics (56). Since, carvacrol and thymol represent the two main active components of oregano oil, it was also included as a test compound and displayed excellent efflux inhibitory activity (57).

Bacterial persisters exhibit a state of dormancy characterized by reduced cellular metabolism and low proton motive force (42). Compounds that modulate the PMF of dormant persister cells can act as attractive leads for anti-persister drug discovery. Moreover, since persisters are known to exhibit enhanced efflux and most efflux pumps being PMF driven, compounds that dissipate membrane potential could be useful in adjunct therapies (58). Hence, fluorescent stain DiBAC_4_(3) was used to evaluate the effect of test compounds on *A. baumannii* membrane potential. Thymol and clove oil showed maximum membrane depolarization (Fig. S4E). The impressive efflux inhibitory activity of thymol, as observed previously could also be attributed to its ability to dissipate *A. baumannii* membrane potential. The results from above assays were compiled and thymol was observed to outcompete other molecules, followed by carvacrol and eugenol (Fig. 4). These compounds were therefore taken up for further validation in *A. baumannii* persister killing assays.

**FIG 4.**
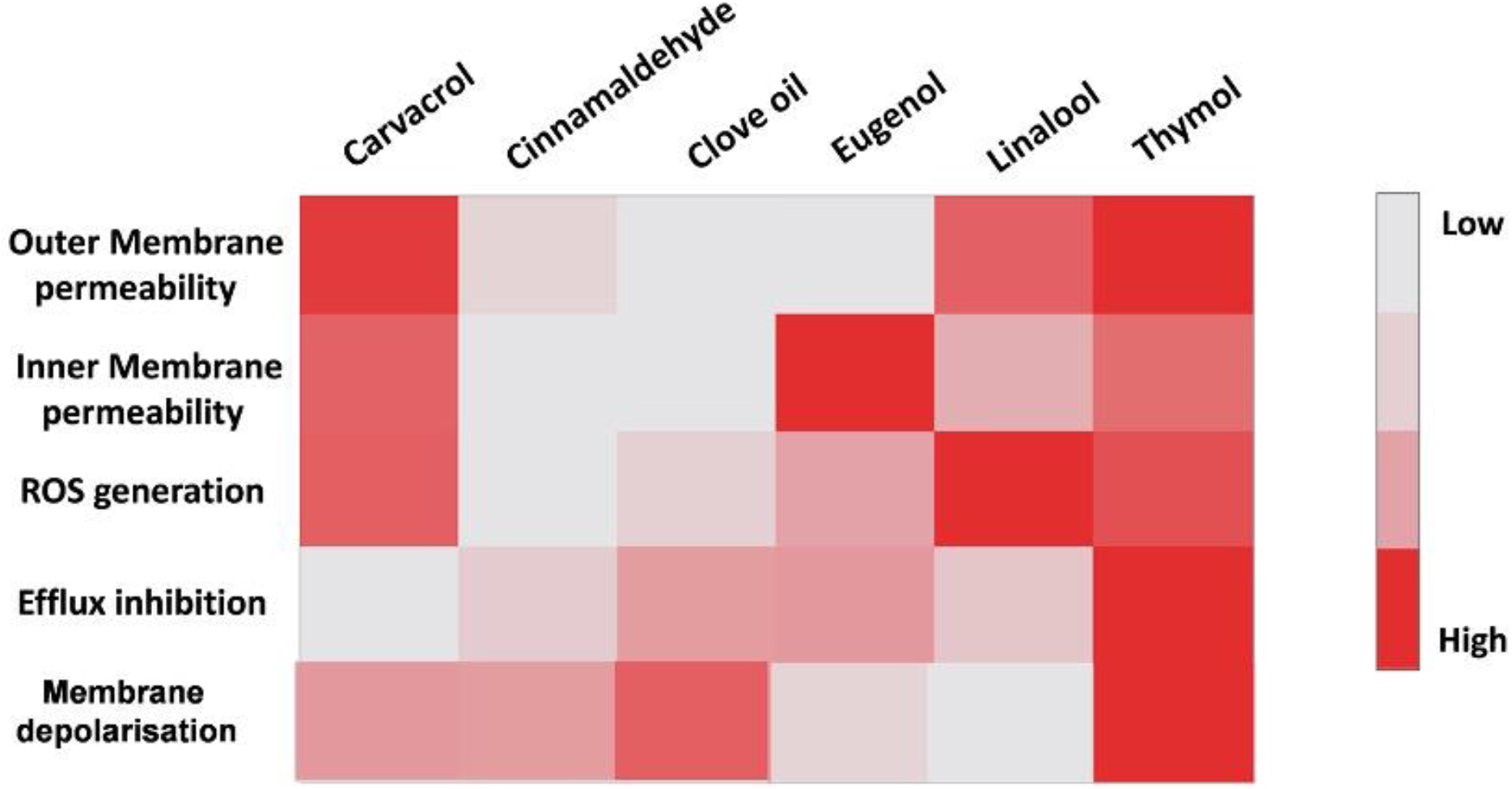
Heat map summarising the results of the mechanistic based screening of GRAS status small molecules against *A. baumannii* AYE. Thymol was chosen as the lead molecule for anti-persister assays.

### Thymol exhibits excellent inhibitory activity against meropenem persisters of *A. baumannii* both in planktonic phase and biofilms

Killing assays against planktonic phase *A. baumannii* persisters induced with bactericidal concentration of meropenem were carried out to validate the anti-persister potential of the compounds screened. Upon exposure to meropenem (100X MIC), cells were incubated with lead compounds for 12h, both in the absence and presence of meropenem. The number of viable bacteria that survived the treatment was evaluated by determining the CFU/mL in comparison to untreated control. As predicted from the results of mechanistic screen, thymol alone was observed to display complete eradication of meropenem induced *A. baumannii* persisters at 1X MIC (Fig. 5A). Interestingly, in combination with meropenem, thymol completely eradicated persisters at lower concentration (0.5X MIC). Eugenol and carvacrol also inhibited meropenem persisters, albeit at higher concentrations (Fig. S5A and S5B).

**FIG 5.**
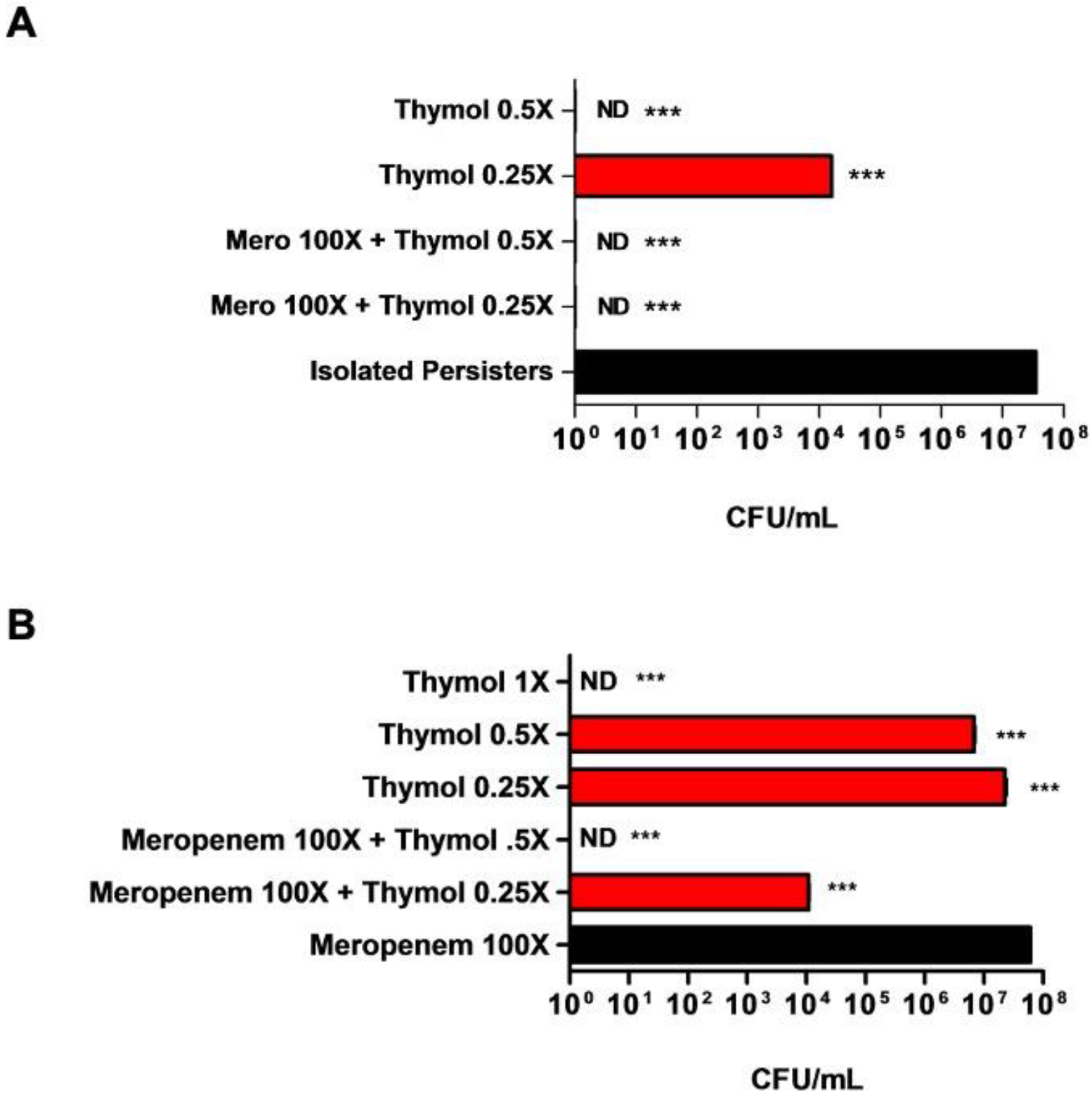
Thymol exhibits excellent inhibitory activity against meropenem persisters of *A. baumannii* AYE both in (A) planktonic phase and (B) biofilms. Each value represents the mean of three values and error bars indicate standard error. One-way ANOVA followed by Tukey’s multiple comparison test was performed (*, P<0.05; **, P<0.01; ***, P<0.0001). ND = not detected.

Recent studies have revealed that bacterial biofilms possess 100–1,000 fold more persister populations than planktonic cultures (59). Hence, the inhibitory activity of compounds against biofilm associated *A. baumannii* persisters was assessed. 48h old *A. baumannii* AYE biofilms were exposed to meropenem alone (100X MIC), compounds alone (at 0.25-2X MIC) or combination of the two. After 12h of respective treatments, number of viable colonies were enumerated. Thymol displayed best inhibitory activity completely eradicating all biofilm associated bacteria, at concentration corresponding to 1X MIC (Fig. 5B). Moreover, thymol-meropenem combination caused complete elimination at 0.5X MIC, thus establishing that co-therapy outcompeted monotherapy, both in case of planktonic and biofilm associated persisters. Eugenol and carvacrol, on the other hand, eradicated biofilm persisters at higher concentrations (Fig. S6A and S6B). It was interesting to note that thymol showed better anti-persister activity to carvacrol, although both are structural isomers (60). This could be attributed to enhanced membrane depolarization and efflux inhibitory potential of thymol, as evident from screening assay.

On the basis of above observations, where meropenem-thymol combination displayed inhibitory potential against *A. baumannii* persisters, we were encouraged to determine if the combination would show a synergistic interaction in a checkerboard assay (61). Interestingly, no synergy was observed with the interaction being additive in nature (FICI ≥ 0.5). This indicated that thymol targeted the mechanisms of antibiotic induced *A.baumannii* persistence under stationary phase conditions.

### Thymol exhibits anti-persister activity irrespective of time of addition

The time of administration of thymol against meropenem induced persistence was further determined by kill kinetics assay, where thymol was added at varying time points (t = 0h, 3h and 6h) during meropenem treatment (100X MIC) of *A. baumannii* AYE SP cells. Complete eradication of *A. baumannii* was achieved within 1h after addition of thymol, irrespective of the time of its administration (Fig. 6).

**FIG 6.**
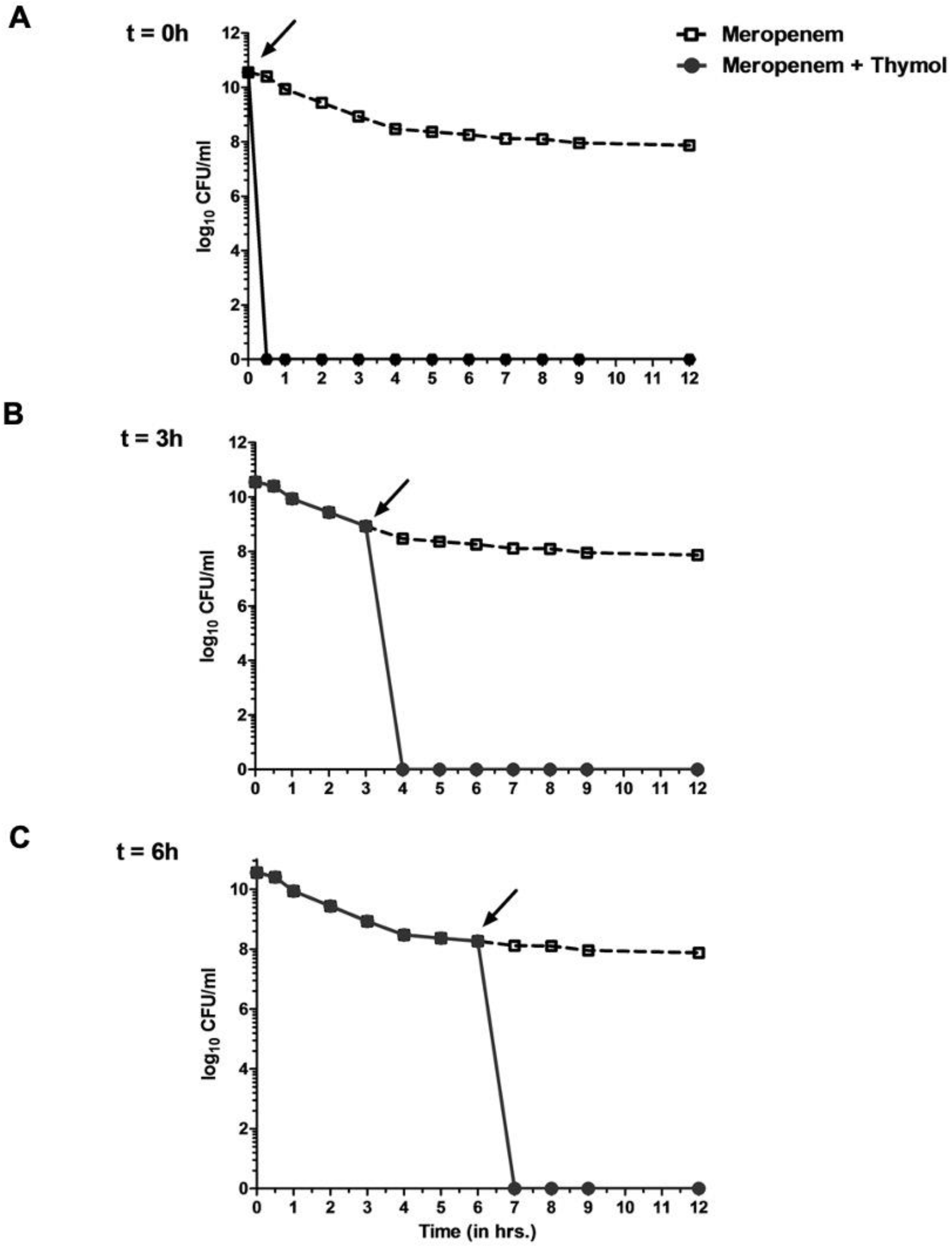
Anti-persister activity of thymol is time-independent. Stationary phase cells of *A. baumannii* AYE were treated with meropenem and thymol was added at time points (A) t = 0h, (B) 3h and (C) 6h as indicated by arrows. Each value represents the mean of three values and error bars indicate standard error. P values were determined by one-way ANOVA followed by Turkey’s multiple comparison test (*, P<0.05; **, P<0.01; ***, P<0.0001).

### Anti-persister activity of thymol is antibiotic independent

We further sought to assess if the anti-persister activity of thymol depended on the nature of the antibiotics that induce persistence. Persisters of rifampicin, tigecycline and polymyxin B were isolated and treated with thymol, both as monotherapy and co-therapy. As shown in Fig. 7A and B, thymol completely eradicated rifampicin and tigecycline induced persisters at 0.25X MIC and 0.5X MIC in combination with antibiotics respectively. On the other hand, polymyxin B induced persisters were completely eradicated by thymol in combination with antibiotic, at 1X MIC (Fig. 7C). These results clearly show that thymol could be combined with different antibiotic classes and thus possess antibiotic independent anti-persister activity.

**FIG 7.**
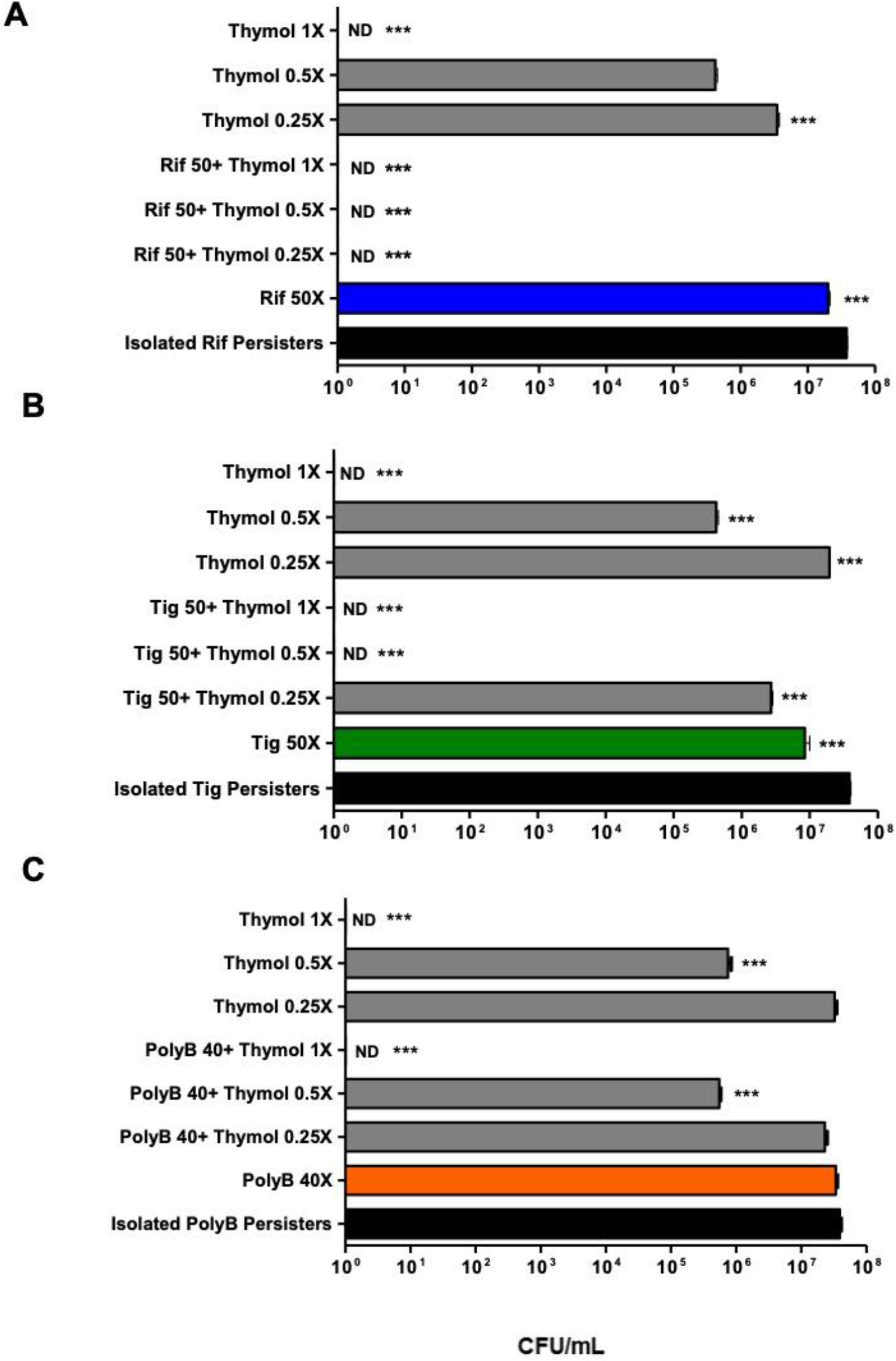
Anti-persister activity of thymol is antibiotic dependent. Thymol inhibits *A. baumannii* AYE persisters of (A) Rifampicin (B) Tigecycline (C) Polymyxin B induced persister cells. Each value represents the mean of three values and error bars indicate standard error. P values were determined by one-way ANOVA followed by Tukey’s multiple comparison test *, P<0.05; **, P<0.01; ***, P<0.0001). ND= not detected

### Thymol exhibits no propensity for resistance generation against *A. baumannii*

The ability of thymol to cause the emergence of resistance in *A. baumannii* AYE was assessed, using the large inoculum approach where 10^10^ CFU/mL of *A. baumannii* were exposed to thymol at 1X and 2X MIC. Thymol displayed no propensity towards resistance development since no colonies were obtained even after 72h of exposure (Table 1).

**Table 1:** Frequency of resistance generation against thymol in *A. baumannii* AYE

| **Frequency of Resistance (FOR)** |  |  |
| --- | --- | --- |
| **Thymol (1X MIC)** | **Thymol (2X MIC)** | **Rifampicin (2X MIC)** |
| $<<< 10^{-10}$ | $<<<10^{-10}$ | $1.2 \times 10^{-3}$ |

### Thymol can inhibit stationary phase cells of *A. baumannii* AYE

The number of persisters in a growing population is known to vary depending on growth phase, with the highest persister frequency observed at stationary phase (62). The bactericidal potential of thymol on stationary phase persisters of *A. baumannii*, cells was evaluated. Treatment of the SP cells with 1X or 2X MIC of thymol alone significantly decreased the number of surviving cells compared to that of the untreated control (Fig. S7A). Scanning electron microscopy of SP *A. baumannii* cells treated with thymol (2X MIC) displayed complete eradication of persisters (Fig. S7B). Thus, thymol exhibits extreme capability of killing both antibiotic induced and stationary phase *A. baumannii* persisters. None the less, thymol inhibited antibiotic induced *A. baumannii* persisters, irrespective of their method of isolation (data not shown).

### Thymol causes membrane depolarization, inhibits efflux and enhances membrane permeability in meropenem persisters

In order to validate the mechanism of persister killing by thymol, we first sought to study its effect on persister membrane potential. *A. baumannii* AYE persisters displayed significant membrane depolarization in response to increasing concentrations of thymol (Fig. 8A). Our study showed meropenem persisters to display reduced permeability and survive exposure to bactericidal meropenem concentrations. Hence, we further performed EtBr efflux assay in isolated persisters in presence of thymol and relative increase in EtBr fluorescence w.r.t. control (no drug) was calculated. As shown in Fig. 8B, thymol caused a dose dependent increase in EtBr fluorescence in *A. baumannii* persisters, thus indicating efflux inhibition. This observation was further confirmed by incubating meropenem persisters with BOCILLIN FL in the presence of thymol which displayed increased fluorescence in comparison to untreated persister cells (Fig. 8C).

**FIG 8.**
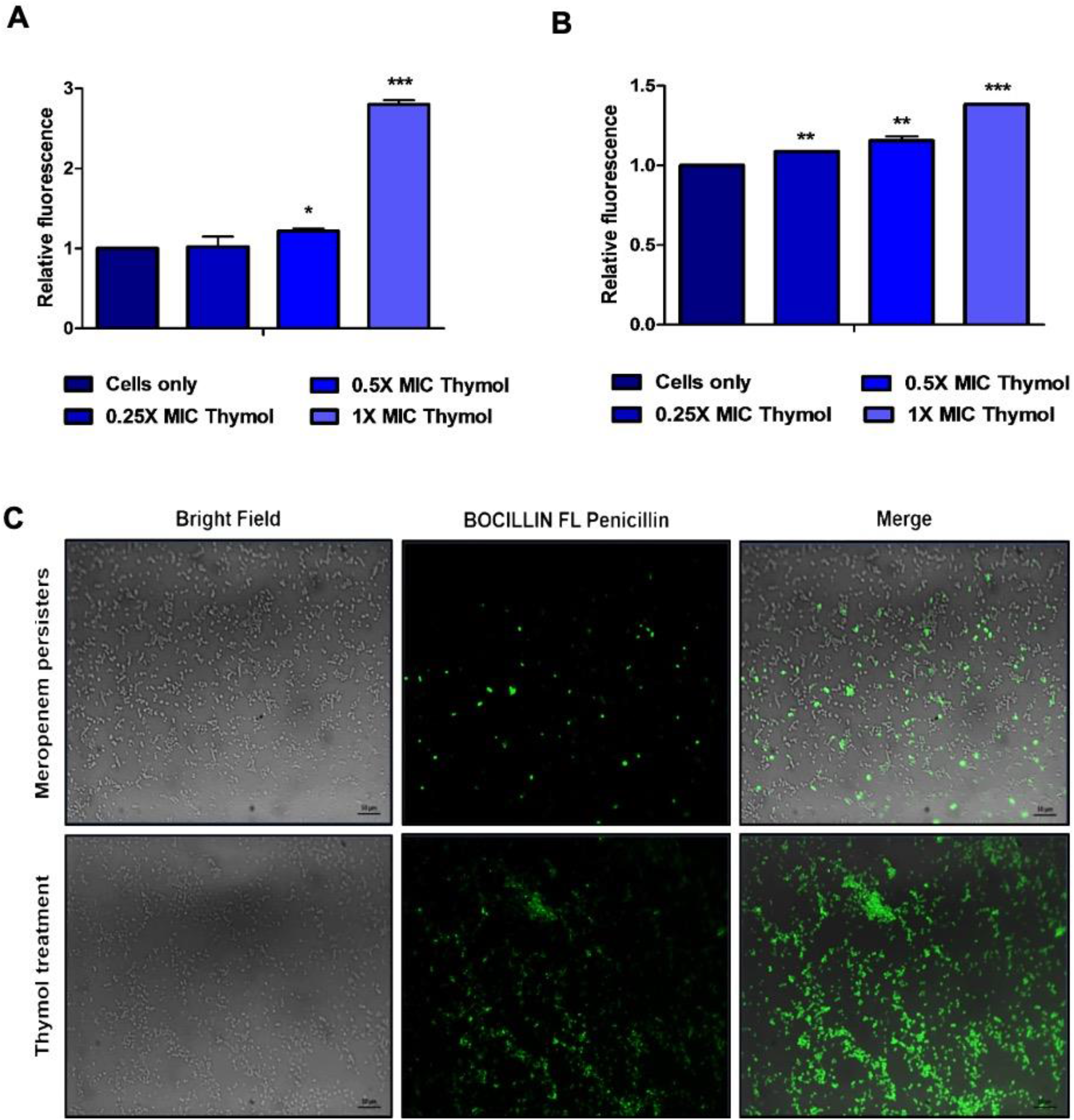
Thymol causes membrane depolarization, inhibits efflux and enhances membrane permeability in meropenem persisters (A) Effect of thymol on membrane potential of meropenem persisters of *A. baumannii* AYE. Relative increase in DiBAC_4_(3) fluorescence in presence of thymol w.r.t. untreated control cells was calculated. (B) EtBr efflux assay in meropenem persisters. Relative increase in EtBr fluorescence upon thymol addition w.r.t. no thymol control was calculated. Data are means±SEM. P values were determined by one-way ANOVA followed by Tukey’s multiple comparison test (*, P<0.05; **, P<0.01; ***, P<0.0001). (C) Fluorescence microscopy to study the effect of thymol on BOCILLIN accumulation in meropenem persisters, in comparison to stationary phase cells. Corresponding brightfield images are shown. Scale bar represents 10 μm.

### Thymol inhibits the respiratory activity of meropenem persisters

As shown previously, the meropenem induced cultures of *A. baumannii* comprised of both persister and VBNC populations. The induction of VBNC state in bacterial pathogen poses a serious health threat and discovery of novel strategies that can induce their resuscitation or inhibit them is of utmost importance (63). Hence, we further sought to assess if thymol could inhibit all viable cells i.e. both persisters and VBNCs, present in the meropenem treated *A. baumannii* cultures, by use of XTT assay and live-dead staining. Encouragingly, thymol showed a significant inhibition in metabolic activity of meropenem persisters at 1X MIC (Fig. 9A). The above observation was further validated by live-dead assay using FM4-64FX in combination with SYTOX green, which is commonly used to assess cell viability (64). As shown in Fig. 9B, thymol inhibited all cells in the meropenem induced persister fractions of *A. baumannii.* Few cells in the isolated persister fractions were observed to stain green (indicating compromised membranes). These represented the meropenem susceptible cells that remained in the isolated fractions, even after the washing steps.

**FIG 9.**
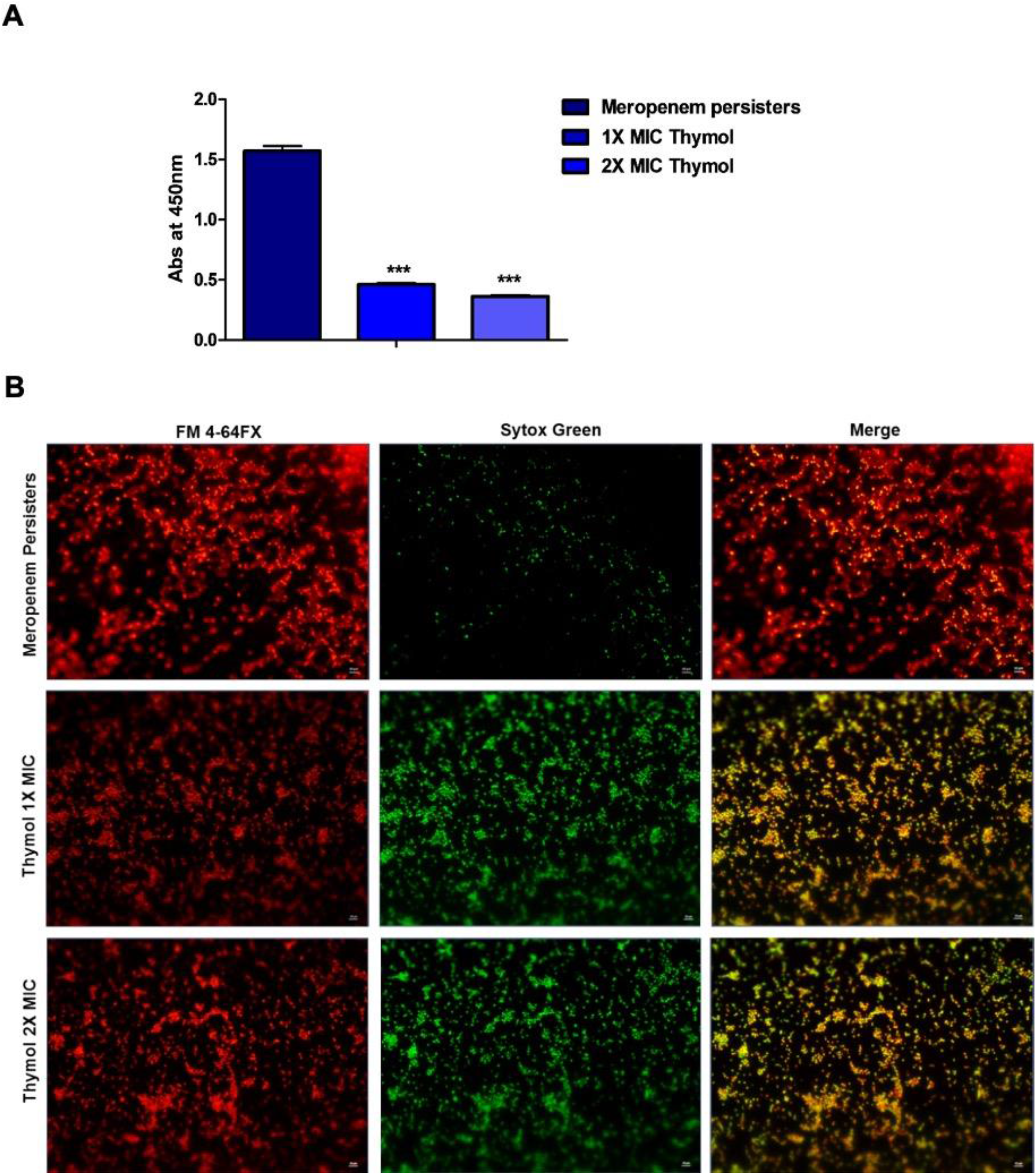
(A) Thymol inhibits the metabolic activity of meropenem induced *A. baumannii* AYE persisters in XTT assay. Each value represents the mean of three values and error bars indicate standard error. P values were determined by one-way ANOVA followed by Turkey’s multiple comparison test (*, P<0.05; **, P<0.01; ***, P<0.0001). (B) Thymol inhibits both culturable and VBNC bacteria present in meropenem induced persister fractions. Viability was assessed using combination of fluorescent dyes FM 4-64 FX and Sytox green. Scale bar represents 10 μm.

The effect of thymol on respiratory activity of *A. baumannii* cells in the stationary phase was also evaluated by XTT assay and live-dead assay. Thymol was observed to significantly inhibit the metabolic activity of SP of *A. baumannii* AYE (Fig.S8A). Live-dead staining of thymol treated *A. baumannii* cells revealed complete killing of the bacterial population at 2X MIC, while partial killing was observed at 1X MIC (Fig.S8B). These observations were found to be well in accordance with the results obtained from the CFU studies for assessing viability counts (Fig. S8A). Since the PMF in bacterial cells is generated by the electron transport chain (ETC), compounds that collapse the PMF can impair electron transport across the respiratory chain, also interfering with the ATP homeostasis. Hence, the ability of thymol to inhibit the respiratory activity of *A. baumannii* persisters could be attributed to its potential to dissipate the PMF.

### Activity of thymol against clinical isolates of *A. baumannii*

The findings obtained so far were further validated by evaluating the activity of thymol against a collection of in-house MDR isolates of *A. baumannii* (Table S2). Stationary phase cultures of *A. baumannii* clinical strains were exposed to meropenem (100-200 μg/ml) for 12h and surviving persister fractions were isolated. All clinical strains displayed a high frequency of persister formation against meropenem. Thymol treatment significantly inhibited meropenem persisters of MDR *A. baumannii* isolates, both as monotherapy and co-therapy (Fig. 10).

**FIG 10.**
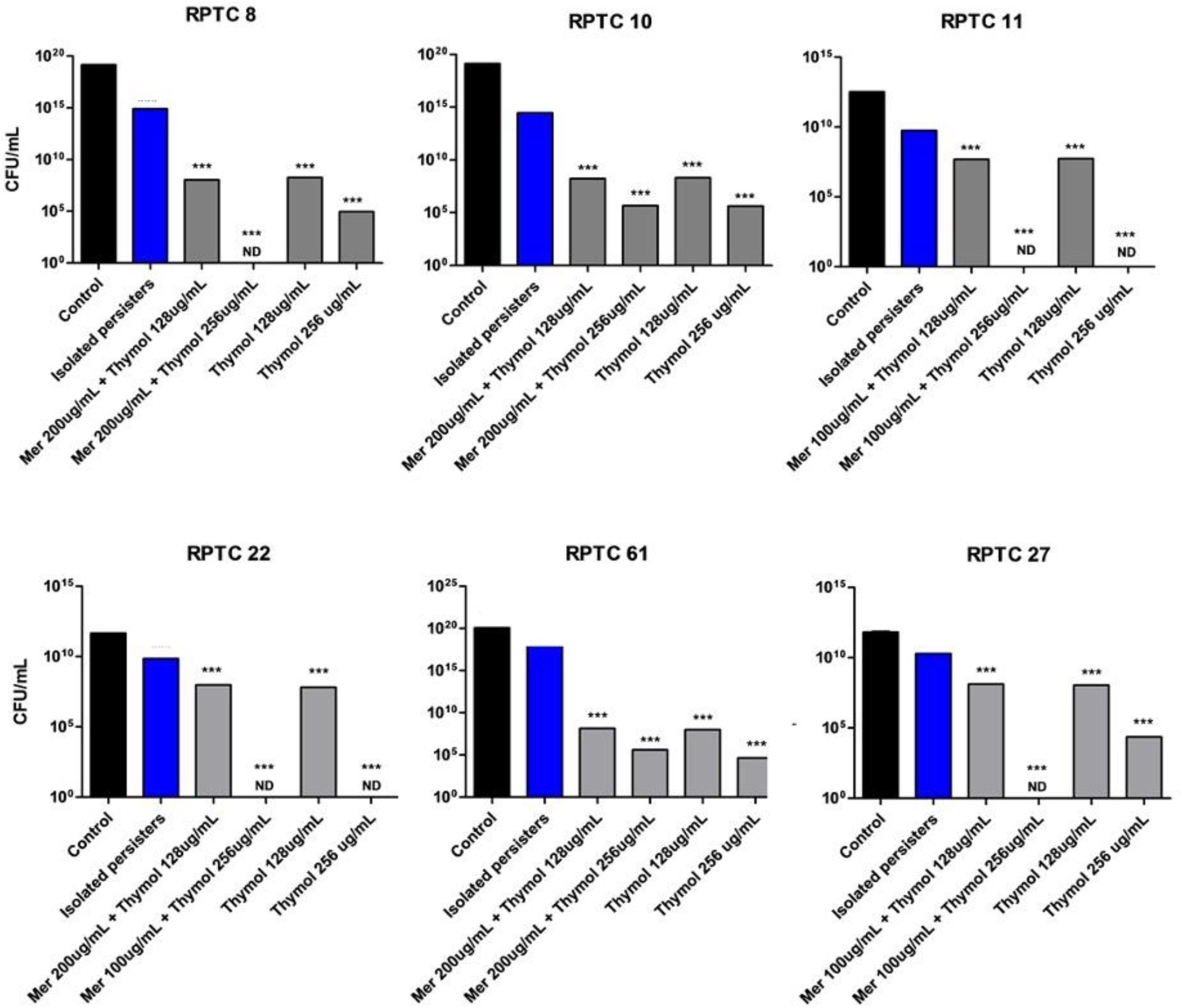
Thymol exhibits anti-persister activity against *A. baumannii* clinical isolates. Isolated meropenem persisters were exposed to thymol at indicated concentrations and viable counts were determined. Each value represents the mean of three values and error bars indicate standard error P values were determined by one-way ANOVA followed by Tukey’s multiple comparison test (*, P<0.05; **, P<0.01; ***, P<0.0001). ND = not detected

### Thymol exhibits anti-persister activity against *K. pneumoniae* and *P. aeruginosa*

Activity of thymol was assessed against other ESKAPE pathogens, namely *K. pneumoniae*, *P. aeruginosa* and *Staphylococcus aureus.* Antibiotic treatment failure of *K. pneumoniae,* a Gram-negative opportunistic pathogen, has been attributed to the presence of both resistant and tolerant strains in the clinical settings (53). Meropenem induced persisters of *K. pneumoniae* ATCC 700698 were isolated and treated with thymol. Complete eradication of *K. pneumoniae* persisters was observed at concentrations as low as 0.25X MIC (i.e. 0.5 mg/mL) of thymol (Fig. 11A). *P. aeruginosa,* another Gram-negative pathogen is known to comprise of antibiotic tolerant subpopulations in the lungs of cystic fibrosis patents. Complete eradication of the meropenem persisters of *P. aeruginosa* was observed upon thymol treatment at 0.25X MIC (i.e. 2 mg/mL) (Fig. 11 B). Vancomycin is the most commonly prescribed antibiotic against severe Methicillin Resistant *S. aureus* infections (65). However, the emergence of vancomycin tolerant strains presents a major clinical challenge. Vancomycin tolerant cells of *S. aureus* ATCC 29213 were isolated and treated with thymol (at 1X MIC i.e. 128 μg/mL). However, thymol did not inhibit vancomycin induced *S. aureus* persisters (data not shown) and therefore its activity was observed to be targeted towards Gram negative pathogens.

**FIG 11.**
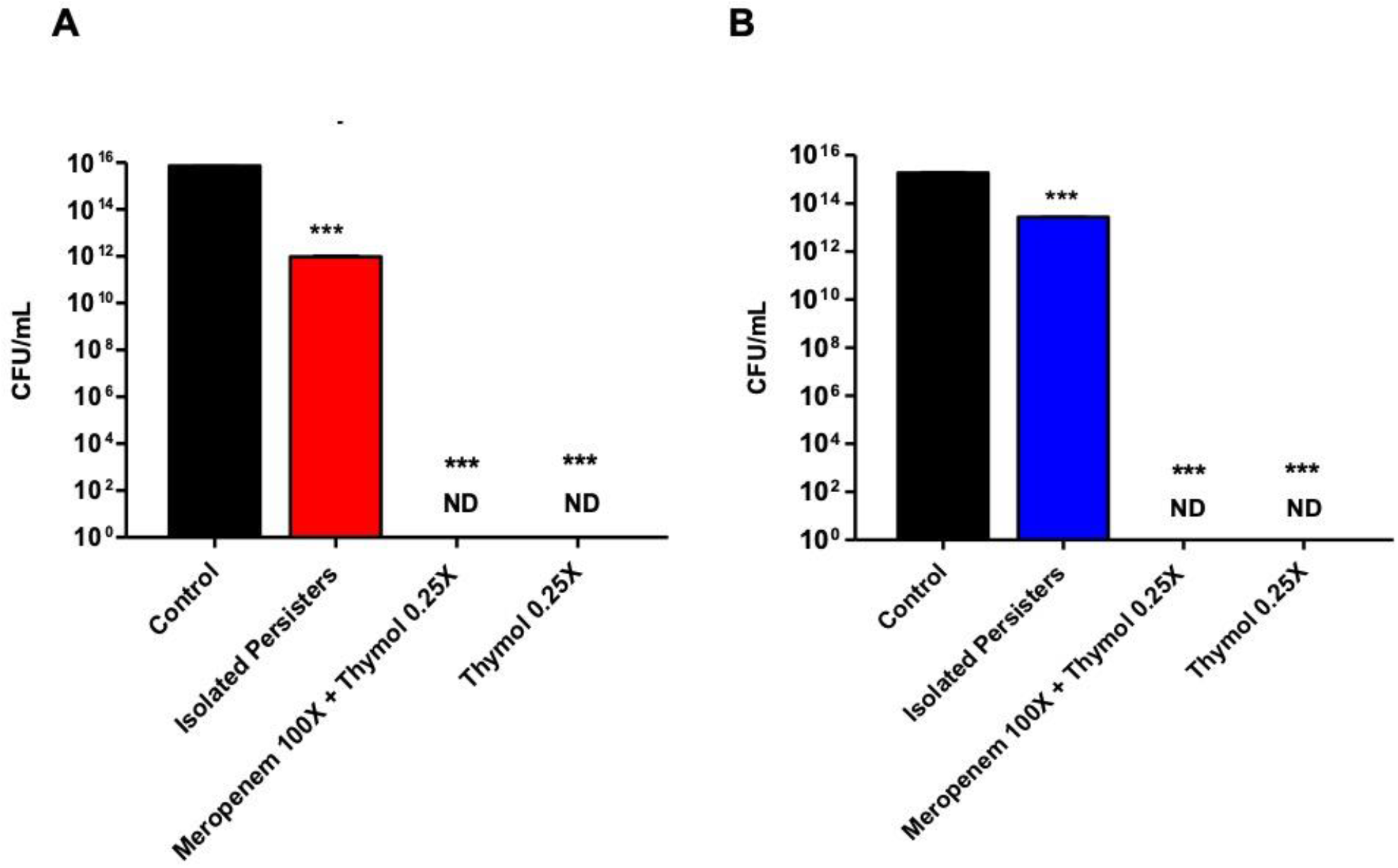
Anti-persister activity of thymol against ESKAPE pathogens (A) *Klebsiella pneumoniae* ATCC 700698 and (B) *Pseudomonas aeruginosa* MTCC 2453. Meropenem induced persisters were isolated and treated with indicated concentrations of thymol, in the absence and presence of meropenem. P values were etermined by one-way ANOVA followed by Tukey’s multiple comparison test (*, P<0.05; **, P<0.01; ***, P<0.0001). ND: not detected

## DISCUSSION

Besides the global problem of multidrug resistance, treatment of infections caused by ESKAPE pathogens is compromised by their ability to give rise to antibiotic-tolerant persister populations (33). Persister cells withstand antibiotic treatment, thereby contributing to the recalcitrance of chronic infections and resistance development (29). In this study, the characteristics and mechanisms of persistence in the ESKAPE pathogen, *A. baumannii* were determined in response to meropenem. Meropenem induced *A. baumannii* persisters exhibited a multi-drug tolerance phenotype, depolarized membranes and decreased permeability. So far, this is the first report that primarily characterizes the mechanisms of meropenem persistence in *A. baumannii.* Similar studies in *S. aureus* show that SP persister formation in response to oxacillin and ciprofloxacin is associated with the occurrence of low membrane potential (66). Although the exact mechanism by which PMF may contribute to *S. aureus* persistence still remains unclear, it was suggested that reduced membrane potential may lead to a halt in cell wall synthesis, as observed in *Bacillus,* thereby causing antibiotic tolerance (66, 67). Meropenen persisters in *A. baumannii* could also follow a similar mechanism, however elaborate studies to delineate the complex persistence mechanisms are very much needed. Recent studies have reported the role of RelA, ppGpp and HigAB, AbkAB toxin/antitoxin (TA) systems towards antibiotic tolerance in *A.baumannii* (68–70). In our study, we decipher the overall role of intracellular ATP levels and membrane permeability in mediating meropenem persistence. Future studies employing a combination of genomics, transcriptomics, proteomics and metabolomics approaches, under conditions best mimicking the host environment of *A. baumannii* infections could be the key for development of new anti-persister treatments.

Our study further identified the anti-persister activity of plant-derived antimicrobial, thymol that could be used as monotherapy or in combination with antibiotics against *A. baumannii* infections. Thymol (2-isopropyl-5-methylphenol) is a monoterpene phenol isolated from plants belonging to the lamiaceae family and can also be chemically synthesized (71). It is used as a food additive and considered to be a potential bioactive compound in the pharmaceutical industry. Thymol has been bestowed the GRAS (generally recognized as safe) status by the Food and Drug Administration (FDA), agency of the United States Department of Health and Human Services (Thymol: 21CFR172.515) (72). In support of its GRAS status, recent studies have reported no associated toxicity in rat models, with LD50 values as high as ~ 4000mg/kg, when administered orally (73). Thymol is long known to possess antibacterial activity against both Gram-negative and Gram-positive pathogens such as *E. coli*, *Salmonella typhimurium*, *Proteus mirabilis* and *Listeria innocua* etc. (74, 75). Only a few studies so far have reported the antibacterial activity of thymol or its derivatives against *A. baumannii* (76). However, none of the studies evaluated the inhibitory potential of thymol against persisters of *A. baumannii* or other ESKAPE pathogens.

Here, we show thymol to possess excellent inhibitory activity against all *A. baumannii* persisters irrespective of their mode of growth (planktonic or biofilms) or culturability. Thymol acted by permeabilising persister membrane, dissipating the membrane potential, triggering ROS production, inhibiting efflux and respiratory activity. It displayed antibiotic independent activity and also inhibited persister populations of *P. aeruginosa* and *K. pneumoniae.* Moreover, it was observed to be refractory to resistance development and could be co-administered during treatment at any time. The ability of thymol to act with mechanistically different classes of antibiotics and its broad spectrum anti-persister activity, indicates thymol to have immense potential to act either alone or as an adjunct in combination therapies against mixed bacterial infections. The anti-persister activity of thymol could further be evaluated in appropriate chronic *in vivo* infection models. Structure activity Relationship (SAR) studies followed by the synthesis of thymol derivatives with improved activity holds immense promise. On the whole, our study encourages future research into the use of thymol for developing novel strategies against chronic and persistent infections.

## MATERIALS AND METHODS

### Bacterial strains and reagents

The strains *A. baumannii* AYE, *A. baumannii* ATCC 17978, *K. pneumoniae* ATCC 700698, *P. aeruginosa* MTCC 2453 and *S. aureus* ATCC 29213 were used for the study. The growth medium was Cation adjusted Muller Hinton Broth (CAMHB), Luria Bertani (LB) broth or LB agar. Antibiotics used in this study were procured from Sigma-Aldrich and Tokyo Chemicals Industries Limited (TCI) and stored as prescribed. Dichloro-dihydro-fluorescein diacetate (DCFH-DA), DiBAC_4_(3), FM 4-64FX, SYTOX green, SYTOX orange and BOCILLIN™ FL Penicillin were purchased from Thermo fisher Scientific. *A. baumannii* clinical isolates were procured from AIIMS, Bhopal and Government Medical College and Hospital, Chandigarh, India.

### Determination of Minimum Inhibitory Concentration (MIC) of antibiotics and GRAS compounds

Serial twofold dilutions of each compound were prepared in CAMHB and added to 96-well plates, as per CLSI recommendations (49). Overnight grown cultures of test strains were sub-cultured in sterile tubes at 37°C and 200 RPM, till they reached OD of 0.5 at 600 nm. Cultures were subsequently diluted to achieve inoculum density of ~10^5^ CFU/ml and added to plates containing the compound dilutions. Plates were incubated for 16 hours at 37°C in a static incubator. At the end of incubation, absorbance was recorded on a plate reader (SpectraMax M2e) at 600 nm.

### Determination of the levels of persistence in *A. baumannii* against meropenem and other antibiotics

Single colony of *A. baumannii* AYE from a freshly streaked plate was inoculated in LB medium for 12h at 37°C and 180 RPM. The culture was 100 times diluted into 5 mL of fresh media and further grown for 12h under similar conditions. Meropenem induced persisters were isolated by exposing 12h old culture of *A. baumannii* AYE (10 mL) to 100X MIC for 12h at 37°C under shaking conditions (26–28). Cultures were then washed thrice with 1X PBS at 4500 g for 4 minutes and resuspended in the same volume of PBS, as of the initial culture. In order to prevent the reversion of isolated persisters into actively growing cells, they were always kept in 1X PBS throughout the assays, with or without meropenem (as indicated in the experiment). Persisters of tigecycline (50X MIC), rifampicin (50X MIC) and polymyxin B (40X MIC) were also isolated following the same procedure. The antibiotic concentrations used for the assay were based on preliminary experiments such that persister levels remained unchanged above the chosen concentrations (except for polymyxin B which demonstrated no surviving fractions above 40X MIC). The frequency of persisters was calculated by dividing the number of cells surviving antibiotic exposure to the initial number of stationary phase cells before treatment. For the meropenem biphasic killing assay, 100X MIC of meropenem was added to the 12h old culture in LB under shaking conditions and the number of surviving bacteria were enumerated at regular intervals by the drop plate method.

Meropenem induced persisters were also isolated by the following procedure, as described previously (24). 12h old culture of *A. baumannii* AYE was diluted 10 times in fresh medium and meropenem (10X MIC) was added for 5 hours. Cells were washed thrice with 1X PBS and spot plated to assess the viable culturable counts. The survival of ATCC 17978 Δ*adeIJK*, Δ*adeIJK*::*adeIJK* and Δ*omp33* strains in the presence of meropenem (10X MIC), in comparison to wild type; was also studied using similar protocol. Expression of AdeIJK efflux pump in ATCC 17978 Δ*adeIJK*::*adeIJK* strain was induced by adding 1mM IPTG to fresh media, at the time of culture dilution (77).

### Meropenem persistence in *A. baumannii* under biofilm conditions

Overnight culture of *A. baumannii* AYE was diluted 100 folds into fresh LB medium and grown until they reached OD_600_ of 0.4. 200 μl cells were added to 96-well plates and biofilms were allowed to form for 48h under static conditions at 37°C. Planktonic cells were removed by washing with PBS followed by meropenem (100X MIC) addition and incubation for 12h. Planktonic cells were washed again and biofilms were disrupted using bath sonicator (at 50 KHz for 10 minutes at RT). The numbers of biofilm associated *A. baumannii* cells that survived meropenem exposure were determined in comparison to untreated control wells.

### Tolerance of meropenem induced *A. baumannii* persisters to other antibiotics

To determine if meropenem induced persisters could exhibit multi-drug tolerance, the survival of isolated meropenem persisters upon exposure to high concentrations of tigecycline (50X MIC), rifampicin (50X MIC) and polymyxin B (20X MIC) was assessed. The numbers of surviving persisters were enumerated after 12h of antibiotic exposure. The fraction of bacteria that exhibit drug tolerance was calculated by dividing the CFU/mL of persisters surviving the test antibiotic exposure to the initial CFU/mL of meropenem persisters, as described previously (78). Similarly, stationary phase *A. baumannii* AYE cells were exposed to test antibiotics respectively and surviving fractions were calculated by determining ratio of cells surviving treatment to the initial number of untreated SP cells.

### Fluorescence Microscopy to assess changes in cellular morphology

Fluorescence microscopy was used to evaluate the morphological changes in *A. baumannii* AYE cells after treatment with bactericidal concentrations of meropenem. *A. baumannii* AYE persister cells were isolated as per the methods described above and was compared with that of untreated control cells. FM 4-64FX (1 μg/ml) was added to the control as well as persister cells and incubated for 15 minutes at room temperature. 10 μL of cells was added to the agar pad (made using 1 % agarose), covered with a cover slip and observed under AXIO A1 ZEISS fluorescence microscope.

### Assay for Outer Membrane Permeability

N-phenyl-1-napthylamine (NPN) was used to assess the outer membrane permeability in the presence of compounds. NPN binds to the outer leaflet of the cell membrane and fluoresces when in the hydrophobic environment. Overnight grown cultures of *A. baumannii* AYE were sub-cultured in fresh medium and incubated at 37°C and 180 RPM up to OD_600_ of 0.5. Cells were washed and resuspended in 5mM HEPES buffer to OD_600_ = 0.3 and incubated with 0.25X MIC of the compounds for 30 minutes in half area black plate (Corning). NPN (10 μM) was added to the wells and fluorescence was immediately measured at an excitation/emission wavelengths of 350/420 nm respectively using a plate reader (SpectraMax M2e). Relative Fluorescence Units (RFU) was calculated by dividing the fluorescence values obtained by the absorbance at 600 nm. Relative increase in fluorescence w.r.t untreated control was plotted.

### Assay for Inner Membrane Permeability

The membrane permeability assay was performed using SYTOX Orange nucleic acid stain which fluoresces upon binding to DNA. *A. baumannii* AYE cells grown up to OD_600_= 0.5. were washed and resuspended in 1X PBS to an optical density of 0.3. SYTOX Orange (1 μM) was added to the cells with constant stirring to let the dye stabilize. Compounds (at 0.25X MIC) and dye loaded cells were added to the 96-well opaque half area plates. Fluorescence was measured after 30 minutes at excitation/emission wavelengths of 488/570 nm respectively and RFU was calculated. Relative fluorescence w.r.t untreated control was plotted.

### Reactive Oxygen Species (ROS) Assay

Exponential phase *A. baumannii* AYE cells (OD_600_ ~ 0.5) were washed and resuspended in 1X PBS to optical density of 0.3 and incubated with Dichloro-dihydro-fluorescein diacetate (DCFH-DA) at 10 μM for 30 minutes at 37°C. Dye treated cells were incubated with compounds (at 0.25X MIC) in 96-well half area black plate (Corning) at 37°C for 2h and read thereafter at excitation/ emission wavelength of 485/528 nm on plate reader (SpectraMax M2e). Relative fluorescence w.r.t untreated control was plotted.

### Ethidium Bromide Efflux Assay

Exponential phase *A. baumannii* AYE cells (OD_600_ ~ 0.5) were washed and resuspended in 1X PBS to optical density of 0.3 and incubated with ethidium bromide (10 μg/mL) at 37°C for 20 minutes. Cells were added to the wells of a 96-well black plate containing test compounds (at 0.25X MIC) and fluorescence was measured at excitation/emission of 480/610nm. Efflux inhibitory potential of thymol against *A. baumannii* persisters was also measured by similar protocol. Meropenem tolerant *A. baumannii* AYE persisters were incubated with EtBr, followed by exposure to varying concentration of thymol. Relative increase in EtBr fluorescence upon thymol addition w.r.t. no thymol control was calculated and plotted.

### Assay for measurement of membrane potential

Exponential phase *A. baumannii* AYE cells were washed and diluted to OD_600_ ~ 0.3 in 1X PBS. Cells were incubated with DiBAC_4_(3) at 10 μM for 30 minutes and three subsequent washing steps with 1X PBS were performed. Cells were added to the wells of a 96-well black plate containing test compounds (at 0.25X MIC) and fluorescence was recorded at excitation-emission wavelength of 490/516 nm in 96-well Corning half area opaque plates. RFU was calculated and relative increase in fluorescence upon thymol addition w.r.t. no thymol control was plotted. In a separate assay, membrane potential of, *A. baumannii* AYE SP cells and isolated meropenem persisters was measured in the presence and absence of thymol (at 1X and 2X MIC).

### Treatment of antibiotic induced persisters with potent anti-persister compounds

Antibiotic induced persisters for meropenem, tigecycline, rifampicin and polymyxin B were isolated as described previously. Isolated persister fractions were washed and resuspended in PBS, divided into 1 ml aliquots and treated with potent GRAS compounds both alone (at 0.25X, 0.5X, 1X MIC and 2X MIC) and in the presence of respective antibiotics for 12h. The numbers of surviving bacteria after compound treatment were enumerated.

### Scanning Electron Microscopy to assess changes in cellular morphology

SEM was performed to evaluate the morphological changes in stationary phase *A. baumannii* AYE cells after treatment with thymol. For sample preparation, *A. baumannii* (12h old culture) was exposed to thymol (1X and 2X MIC) for 12h. Treated cells were washed thrice with 1X PBS and centrifuged at 8000g for 4 min and fixed overnight at 4°C with 2.5% glutaraldehyde. Fixed cells were further dehydrated with an ethanol gradient, coated with a layer of gold and visualized under SEM.

### Checkerboard assay

Interactions between antibacterial compounds can be assessed using fractional inhibitory index (FIC) values calculated from checkerboard assays (79). 100 μl of LB broth was distributed into each well of 96-well the microtitre plates and meropenem was serially two-fold diluted along its ordinate, while thymol was diluted along the abscissa. 100 μL of *A. baumannii* AYE cells (~ 10^5^ CFU/ml) in LB medium were added to the plates and incubated at 37°C. Absorbance was measured after 16h. The ΣFICs were calculated as follows: FICI = FIC A + FIC B, where FIC A is the MIC of drug A in the combination/MIC of drug A alone, and FIC B is the MIC of drug B in the combination/MIC of drug B alone. The combination is considered synergistic when the FICI is ≤0.5, indifferent/additive when the FICI is >0.5 to <2, and antagonistic when the FICI is ≥2.

### Killing assays to assess anti-persister potential of GRAS compounds

In order to assess the anti-persister potential of test compounds against *A. baumannii* AYE in planktonic phase, SP cells or meropenem persisters were incubated with varying concentration of test compounds (0.25X, 0.5X, 1X and 2X MIC) with or without meropenem (100X MIC), under shaking conditions. The number of viable bacteria was enumerated after 12h of incubation. For killing assays under biofilm conditions, 48h old *A. baumannii* AYE biofilms were treated in 1X PBS with meropenem alone (at 100X MIC), GRAS compounds (at 0.25X MIC, 0.5X MIC, IX MIC and 2X MIC) or a combination of both. After 12h of incubation under static condition at 37°C, biofilm associated cells were dislodged and plated to enumerate the number of viable colonies.

### Assay to determine killing kinetics of thymol

The kinetics of inhibition of *A. baumannii* cells in the presence of meropenem (at 100X MIC) upon addition of thymol at different time intervals was studied, as described previously (80). SP cells of *A. baumannii* AYE were treated with meropenem (100X MIC) or combination of meropenem and 1X MIC of thymol. Thymol was added to the cultures at different time points t = 0h, 3h and 6h. The number of viable cells was determined at regular intervals in terms of CFU/mL.

### Assay for anti-persister potential of thymol against persisters of other antibiotics

Anti-persister activity of thymol against persisters of rifampicin, tigecycline and polymyxin B was assessed using the following procedure. SP cells of *A. baumannii* AYE were exposed to rifampicin (50X MIC), tigecycline (50X MIC) and polymyxin B (40X MIC) for 12h to induce persister formation. Isolated persisters were then treated with thymol at 0.25X, 0.5X and 1X MIC in the absence and presence of respective antibiotics. After 12h of incubation, the cells were plated to enumerate the number of viable colonies.

### Fluorescence microscopy to assess the accumulation of BOCILLIN™ FL Penicillin

Isolated meropenem induced *A. baumannii* AYE persisters in IX PBS were incubated with BOCILLIN™ FL Penicillin (10 μg/ml) for 30 minutes in the presence of meropenem (100X MIC) to maintain persistence. Cells were then washed with IX PBS, deposited on 1% agarose pads and visualised under a Zeiss Axioscope A1 fluorescence microscope equipped with an AxioCam MRC digital camera using EC Plan-Neofluar 100X objective. For comparison, stationary phase cultures of *A. baumannii* AYE were washed, resuspended in 1X PBS, and similarly incubated with BOCILLIN followed by microscopy studies.

### Assay for respiratory activity Tetrazolium salt (XTT) reduction assay

Meropenem induced persisters and SP cells of *A. baumannii* AYE were washed thrice with 1X PBS and resuspended in the same buffer. XTT dye (0.5 mg/ml) in combination with menadione (50 μM) was added to the cells resuspended in 1X PBS and incubated for 2h. Absorbance was read at 450nm to assess the reduction of XTT dye and the formation of soluble formazan. Prior to the experiment, CFU/mL of log phase and SP cells were normalized to the CFU/mL of isolated meropenem persisters.

### Live-Dead staining to assess cell viability

Thymol treated *A. baumannii* AYE persisters were harvested and stained with membrane dye FM™ 4-64FX (1 μg/mL) and nucleoid stain SYTOX green (1 μM) at room temperature for 15 minutes. A small volume (5μl) of cells was deposited on agarose pad and sealed with a clean coverslip. The cells were then observed under a Zeiss Axioscope A1 fluorescence microscope equipped with an AxioCam MRC digital camera using EC Plan-Neofluar 100X objective to assess viability.

### ATP quantification assay

Intracellular ATP levels in *A. baumannii* AYE SP cells and meropenem persisters were determined using the BacTiter-Glo™ Microbial Cell Viability Assay kit (Promega) which determines the number of viable cells based on quantitation of ATP. Persisters were isolated as described previously and culturable CFUs were enumerated by plating serially dilutions. Total number of cells in the persister fractions were determined by staining cells with DAPI and analyzing by flow cytometry. Luminescence units were normalized against both CFUs and cell numbers for ATP quantification.

### Frequency of resistance (FOR) of thymol against *A. baumannii* AYE

A log phase culture of *A. baumannii* AYE (OD_600_ = 0.5) was centrifuged at 7000 RPM for 4 minutes and diluted in fresh LB medium in order to obtain ~ 10^10^ CFU/ml of cells as the starting inoculum. Cells were serially diluted and plated to confirm the initial CFU count. The cell suspension was then divided into 1 mL aliquots and exposed to thymol at 1X and 2X MIC for 72h at 37°C under shaking condition. Rifampicin at 2X MIC served as the positive control for the assay. At the end of incubation, treated and untreated cells were serially diluted and plated. Frequency of Resistance was calculated by dividing the number of CFU obtained on the drug treated cultures by the number of bacteria in the untreated populations.

## ACKNOWLEDGEMENTS

We thank Prof. Ayush Kumar, University of Manitoba, Canada for sharing the strains ATCC 17978 Δ*adeIJK* and ATCC 17978 Δ*adeIJK*::*adeIJK*. The authors are also thankful to Prof. Naveen K. Navani, IIT Roorkee, India for sharing the ATCC 17978 Omp33 mutant. TB was supported by Senior Research fellowship from MHRD, Govt. of India. AC was supported by PG scholarship from DBT, Govt. of India and SRU was supported MHRD post-doctoral fellowship.

